# Multidirectional measures of shear modulus in skeletal muscle

**DOI:** 10.1101/2021.08.01.454699

**Authors:** William E. Reyna, Eric J. Perreault, Daniel Ludvig

**Author notes:** **Corresponding Author:** Daniel Ludvig, Address: Department of Biomedical Engineering, 2145 Sheridan Road, Evanston, IL 60208.

## Abstract

The material properties of muscle play a central role in how muscle resists joint motion, transmits forces internally, and repairs itself. While many studies have evaluated muscle’s tensile material properties, few have investigated muscle’s shear properties. None of which have taken into account muscle’s anisotropic structure or investigated how different muscle architecture affect muscle’s shear properties. The objective of this study was to quantify the shear moduli of skeletal muscle in three orientations relevant to the function of whole muscle. We collected data from the extensor digitorum longus, tibialis anterior, and soleus harvested from both hindlimbs of 12 rats. These muscles were chosen to further evaluate the consistency of shear moduli across muscles with different architectures. We calculated the shear modulus in three orientations: parallel, perpendicular, and across with respect to muscle fiber alignment; while the muscle was subjected to increasing shear strain. For all muscles and orientations, the shear modulus increased with increasing strain. The shear modulus measured perpendicular to fibers was greater than in any other orientation. Despite architectural differences between muscles, we did not find a significant effect of muscle type on shear modulus. Our results show that in rat, muscles’ shear moduli vary with respect to fiber orientation and are not influenced by architectural differences in muscles.

## Introduction

The shear modulus, a value quantifying a material’s resistance to shear deformation, is thought to be the primary factor in load transfer within muscle (Huijing, 1999). These loads can be internal from active or passive muscle function or external from injury (Huijing, 1999; Jarvinen et al., 2013). Most of muscle research has been on the active properties of muscle, as they are most relevant to voluntary movement. However, the passive properties of muscle play a central role in how muscle resists joint motion, transmits forces internally, and repairs (Herbert and Gandevia, 2019). Passive forces on muscle can be parallel, perpendicular, or across muscle fibers. This can result in purely shear or tensile deformation, or a combination of the two. While recent studies have begun to investigate passive stiffness in tension (Bosboom et al., 2001; Lieber and Friden, 2019; Morrow et al., 2010), there remains a void in understanding muscle’s anisotropic shear properties. Therefore, it is necessary to quantify the three-dimensional (3D) shear modulus of muscle. This will allow us to better interpret the anisotropic function of muscle and its resistance to shearing forces.

Muscle has often been described as a transversely isotropic material based on the geometric arrangement of its fibers (Blemker and Delp, 2005). Material properties parallel to the fibers are typically considered to differ from those in the plane of symmetry perpendicular to the fibers. In tension, muscle has two primary directions: 1) longitudinal (parallel) and 2) transverse (perpendicular) to muscle fibers. Any measurement taken perpendicular to muscle fibers will result in the same Young’s modulus. However, in shear, where forces are applied in planes, the muscle have three defining directions: 1) parallel, 2) perpendicular, and 3) across muscle fibers (Fig. 1). For continuum modeling, measuring the shear modulus in these three directions would allow us to describe muscle under purely shear strain or stress. Additionally, these directions encompass the physiological loading directions and those that may occur with injury, such as blunt force trauma. Without these multidirectional measures of shear, we limit our understanding of muscle to forces oriented parallel to fibers, thereby excluding the potential for simulating injuries, or shear wave propagation, with shearing forces in other directions. Furthermore, combining 3D measures of shear modulus with measures of Young’s modulus and Poisson ratio we gain the ability to model muscle under any stress or strain. Currently, to our knowledge, direct measures of a muscle’s shear modulus have been limited to a single dimension. Previous work by Morrow et al. (2010) used a novel apparatus to directly quantify Young’s modulus in two dimensions and shear modulus in one dimension. They made measurements in longitudinal extension, transverse extension, and lateral shear of whole muscle. They concluded that the Young’s modulus in tension was higher in the fiber direction (parallel) than in the cross-fiber (perpendicular) direction. However, as they only measured one direction of shear, the 3D characterization and anisotropy of the shear modulus remains to be determined.

**Figure 1.**
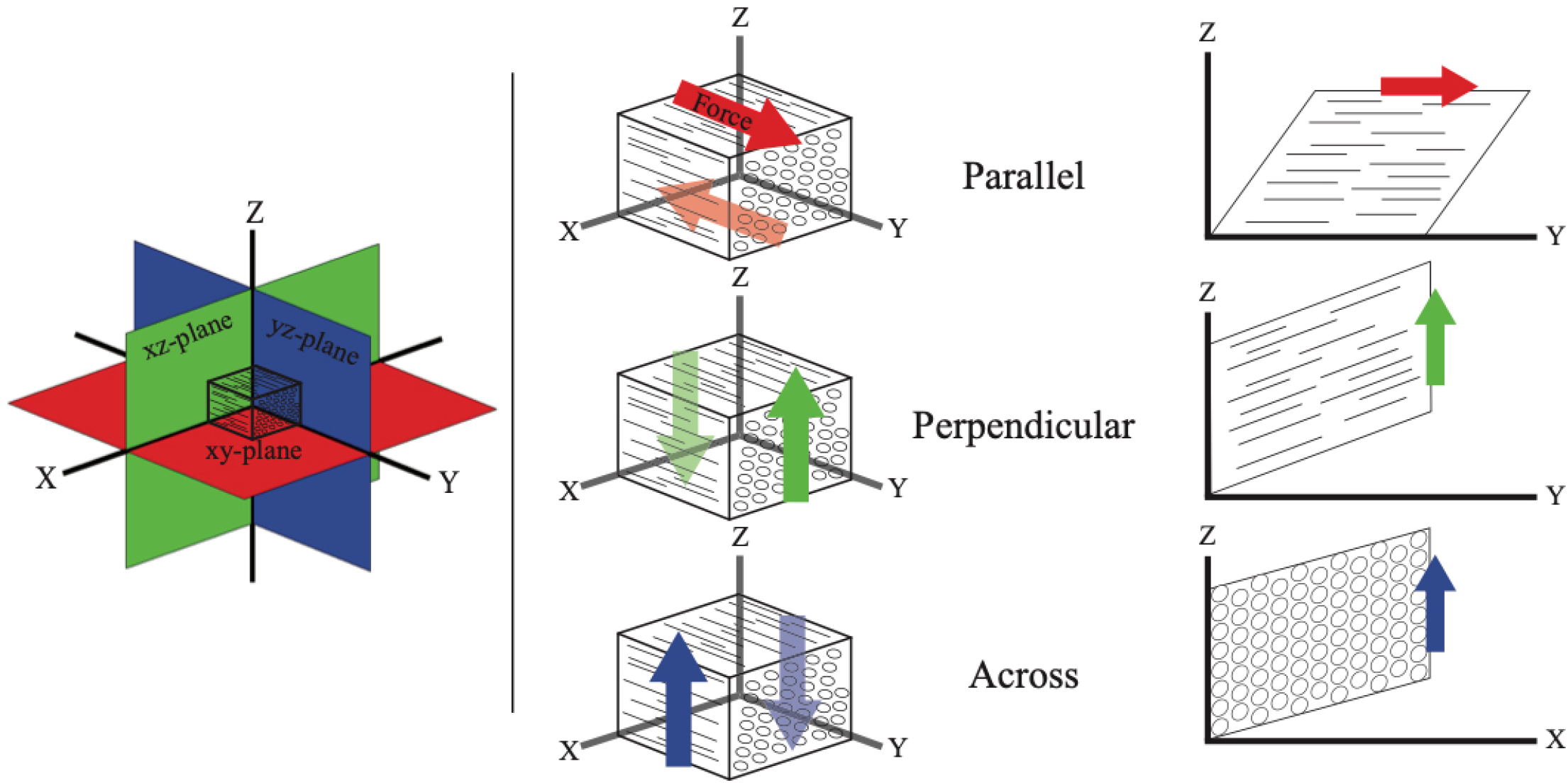
Three-dimensional representation of shearing muscle in the parallel (xy-plane), perpendicular (xz-plane), and across (yz-plane). Solid arrows indicate direction of force and displacement, while transparent arrows indicate stationary resistant force.

It is unknown how differences in muscle architecture alter the shear modulus of muscle. Muscles differ in architectural features like length, cross-sectional area, fiber packing and muscle fiber arrangement (Gans, 1982; Lieber and Friden, 2000). These architectural differences change the amount of excursion and how passive load is distributed through the muscle (Burkholder et al., 1994; Gans, 1982). One of the most striking differences in muscle can be pennation angle, a measure of fiber orientation in reference to the longitudinal axis of muscle. In rat hindlimbs, the pennation angle of muscles can range from around 0 to 20 degrees (Eng et al., 2008). These differences in fiber orientation can result in varying levels of force generation (Lieber and Friden, 2000) and transmission (Huijing, 1999). It has been proposed that for a given change in fiber length, more resistance will be incurred as the pennation angle increases (Huijing, 1999). Therefore, it is important to evaluate the consistency of shear modulus measures across muscles with differing architectures. This will provide further clarity into the potential relationship between shear modulus and muscle architecture.

The objective of this study was to quantify the shear moduli of skeletal muscle in the three orientations relevant to the function of whole muscles. Our primary hypothesis was that the shear modulus would differ with in the different planes of shear relative to the orientation of the muscle fibers. We also hypothesized that the shear modulus would be different across muscle types due to differences in their architectural features. We tested these hypotheses by measuring the shear modulus in the extensor digitorum longus (EDL), tibialis anterior (TA), and soleus (SOL) from rat hindlimbs. Measures were made at three orientations: parallel to, perpendicular to and across the muscle fibers. Our results provide multidimensional estimates of shear moduli that can be used in three-dimensional modeling. This is an important step towards evaluating shearing forces in muscle and further understanding the role of muscle material properties in force transmission and muscle injury.

## Materials and Methods

All data were collected from healthy Sprague-Dawley rats obtained through the Northwestern University tissue sharing program. A total of 12 rats were used. We evaluated three muscles with differing architectures to determine if shear modulus was consistent across muscles. The typical pennation angle in rat of: the EDL is 9.0 ±1.1 degrees; the TA 12.8 ±1.2 degrees; and the SOL is 3.9 ±2.4 (Eng et al., 2008).

### Tissue Preparation

The EDL, TA, and soleus muscles were harvested from both hindlimbs immediately following termination and placed in chilled phosphate-buffered saline to stem the effects of rigor (Tuttle et al., 2014). Muscle specimens had their aponeuroses dissected away using a surgical scalpel to ensure measurements were made only on muscle tissue, similar in procedure to Morrow et al. (2010). Before mechanical testing, muscles were sectioned into rectangular cubes (~9 x 9 x 4 mm) aligned with the muscle fibers. Only a single cube was obtained from each specimen. Shear measurements were made with the cubes oriented in one of three directions: parallel, perpendicular, or across muscle fibers, allowing us to obtain 3D measures of the shear modulus (Fig. 1). Data were collected from 106 total specimens. Specimens (24 total) were eliminated if they showed separation from our testing apparatus during testing or artifacts (drops and rises) were observed in the raw data. The length, width, and thickness of each specimen was measured prior to testing. These physical properties were used in the calculation of stress and strain.

### Mechanical Testing

Shear modulus was measured using an Instron mechanical tester (Instron 5942; Instron Corp., Canton, MA). An offset aluminum bracket with acrylic inserts was mounted into the uniaxial tensile tester, similar in design to Morrow et al. (2010) (Fig. 2A). Aluminum brackets were milled in unison to ensure that surfaces were parallel with one another. Acrylic inserts were used to promote adhesion of the tissue and to easily exchange the testing surface between specimens. The harvested muscle specimens were fixed to the acrylic plates using cyanoacrylate glue. The plates were then screwed onto the aluminum brackets to form a rigid clamp. Specimens were positioned in the center of the testing apparatus to ensure that forces were uniformly applied to the tissue faces and that measurements were in line with the load cell. The use of cyanoacrylate glue bonded tissue to the fixture beyond muscle failure, as seen by rupture within samples as opposed to separation from our mounting plates.

**Figure 2.**
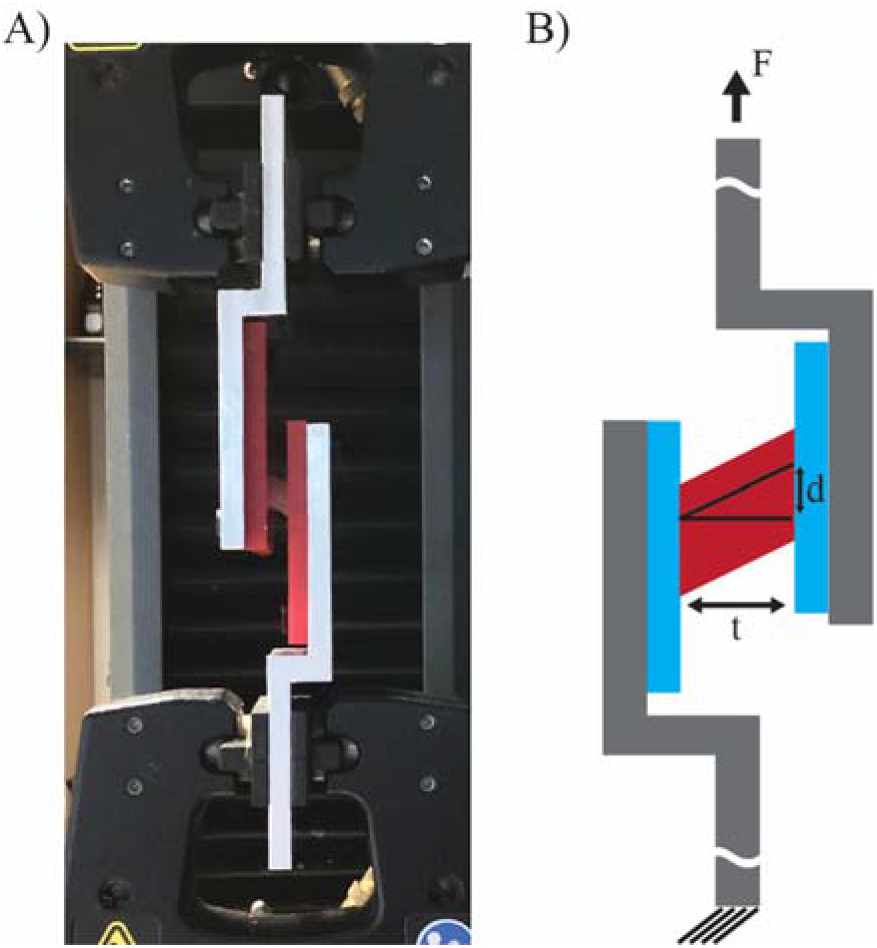
Shear testing apparatus A) photograph and B) schematic. Exaggerated representations of the thickness (t), displacement (d), and force (F) are show in the schematic.

Muscle specimens were tested within 12 hours of harvesting. Data were collected at a strain rate of 5% per minute until failure, as indicated by a sudden drop in force and tearing of the muscle.

### Data Analysis

Shear modulus was calculated from measures of shear stress and shear strain. Tissue shear stress, specifically the second Piola-Kirchhoff stress, was calculated as the applied force (F) divided by the initial cross-sectional area (Fig. 2B). The second Piola-Kirchhoff stress is used as it references the undeformed state of a material with a constant cross-section. Our cubic specimens, to the best of our abilities, have a constant cross-section and our adhered surface encompasses the entire face of the specimen. Tissue shear strain, specifically the Green shear strain, was calculated as the inverse tangent of displacement (d) divided by initial thickness (t) (Fig. 2B). Green shear strain was used because of the small deformations in length. Representative stress-strain data are shown in Fig. 3. Instantaneous shear modulus was computed as the slope of the stress-strain curve (Fig. 3). We report values at regular strain intervals of 0.02, within the predicted physiological range of 0.1 to 0.4 (Blemker et al., 2005).

**Figure 3.**
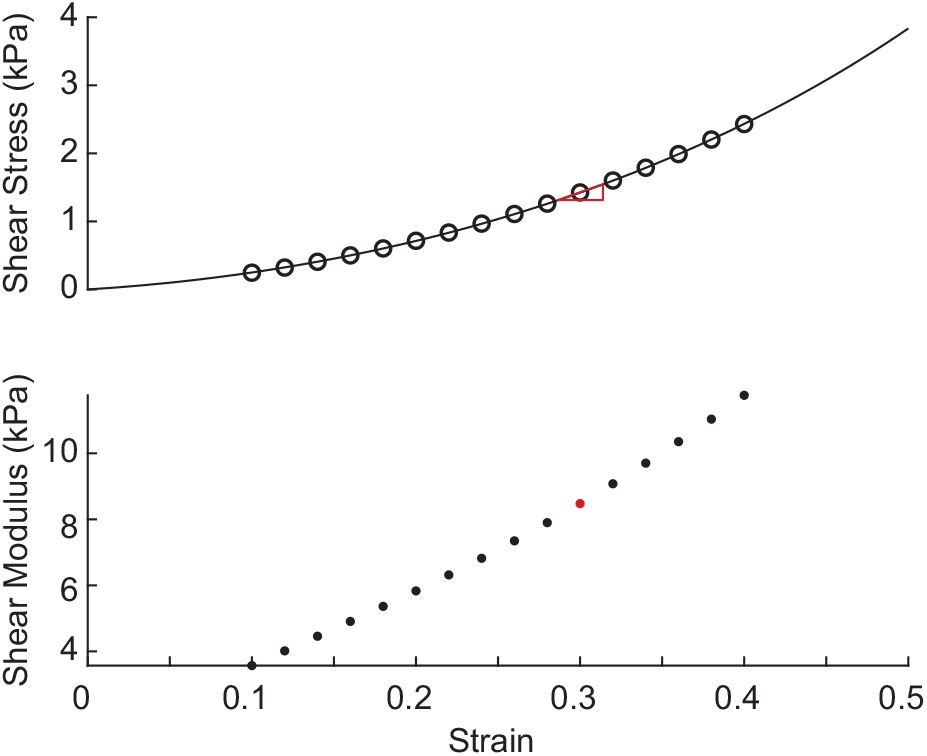
Representative data for shear stress and shear modulus versus strain from a single specimen. Open circles indicate the strain values selected within the physiological strain limit at which the shear modulus was calculated. Shear modulus was calculated as the slope of the shear stress-strain relationship, whose calculation for a single point is denoted by the red triangle.

### Statistical Analyses

We compared the shear modulus across the three different orientations and the three different muscles using a linear mixed-effects model. We fitted a model with the shear modulus as the dependent variable, strain as a continuous factors, muscle and orientation as fixed factors, and specimen as a random factor. Our primary hypothesis was that the shear modulus would differ with the orientation of the shear with respect to the muscle fibers. We tested this hypothesis by using an F-test to determine whether there was a significant effect of orientation or its interactions with strain and muscle on the shear modulus. Our secondary hypothesis was that the shear modulus would differ between the different muscle types. Similarly, we tested this using a F-test to determine if there was a significant effect of muscle type and its interactions with strain and orientation on the shear modulus. For both our primary and secondary hypothesis we ran post-hoc analysis as necessary to determine which orientations and muscles were different from the others. In the post-hoc tests, if we were testing for multiple parameters then a F-statistic was used; if we were testing for only a single parameter then a t-statistic was used. For all statistical tests, a Satterthwaite degrees of freedom approximation was used (Luke, 2017), along with a significance level of α=0.05. All analyses were carried out in Matlab (Mathworks, Natick, MA). Results are presented as mean ± standard error, unless otherwise specified.

## Results

The shear modulus increased with increasing strain on the muscle. This was observed for all muscles and orientations, as seen in the example specimens shown in Fig 4. The increase in shear modulus with strain was fairly linear, as demonstrated by the goodness-of-fit of our linear mixed-effects model (r^2^ = 0.98) for describing the data across all specimens. On average shear modulus increased 8 ±1 kPa (p < 0.001) over a strain of 1. This resulted in a shear modulus that is 2.3 times greater at strain of 0.4 (the top of the physiological range) compared to unstrained state.

**Figure 4.**
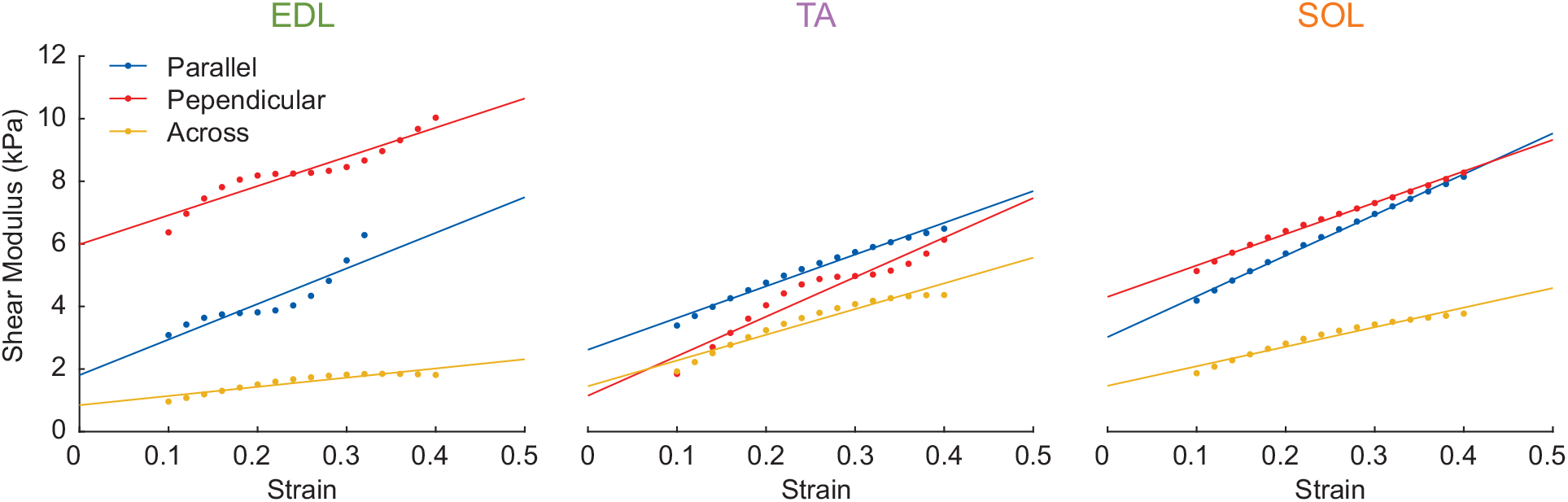
Representative date showing shear modulus as a function of strain for all combinations of muscle and orientation. The fitted line comes from our linear mixed-effects model, which was an extremely good fit for our data overall (R^2^ = 0.98)

The shear modulus differed with the orientation of the muscle (Fig. 5; F_12,73_ = 2.3, p = 0.015). Since the orientation could affect both the shear modulus at zero strain and the slope at which the shear modulus increases with strain, we first tested for a combined effect of orientation on both parameters. We found the shear modulus was different in the perpendicular orientation compared to the parallel (F_2,73_ = 3.8, p = 0.027) or across (F_2,73_ = 8.6, p < 0.001) orientations. The difference between the parallel and across orientations did not reach statistical significance (F_2,73_ = 1.3, p = 0.29). The difference between the perpendicular orientation and the other two orientations was due to a combination of both the increased shear modulus at zero strain and an increased shear modulus-strain slope. The shear modulus at zero strain was greater in the perpendicular orientation (3.9 ± 0.6 kPa) compared to the parallel (2.4 ± 0.4 kPa, t_73_ = 1.5, p = 0.13) or the across (1.5 ± 0.5 kPa, t_73_ = 2.3, p = 0.03) orientation. The slope of the shear modulus-strain relationship was also greater in the perpendicular orientation (11 ± 10 kPa) than the parallel (8 ± 7 kPa, t_73_ = 0.8, p = 0.57) or the across orientations (6 ± 8 kPa, t_73_ = 1.2, p = 0.25). However, due to the large variability in the slope of the shear modulus-strain relationship across the specimens, none of the differences in slope reached statistical significance. The combined effect of increased shear modulus at zero strain, and an increased shear modulus-strain slope resulted in a shear modulus that was significantly greater in the perpendicular orientation than the other orientations at all physiological strain (0.1 – 0.4) values (Fig. 5).

**Figure 5.**
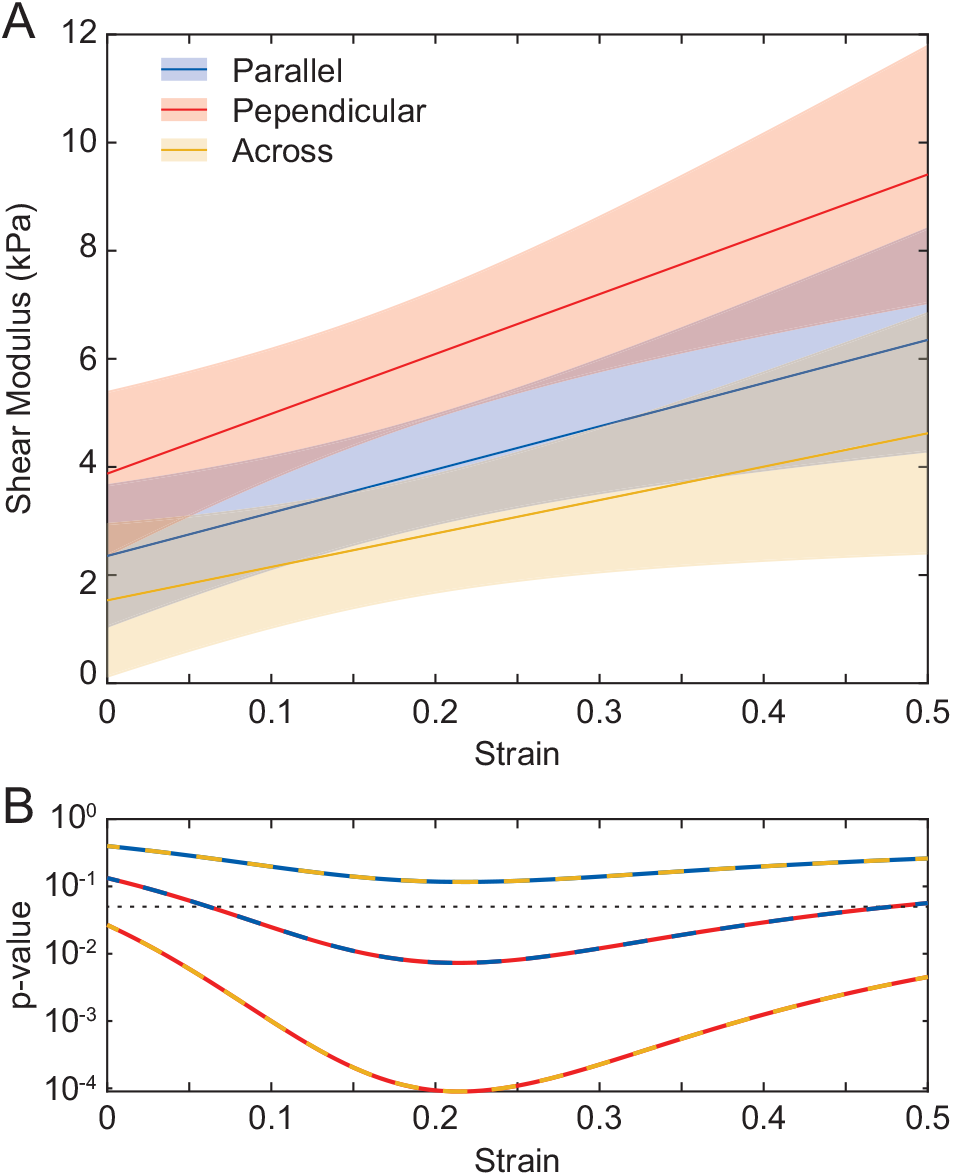
Shear modulus differed with orientation of the muscles. A) Shear modulus was largest when the muscle was oriented perpendicular to the fibers, followed by when the muscle was parallel, and lowest when the muscle was oriented across the fibers. Shaded area represents 95% confidence interval. B) This difference in shear moduli reached the level of significance (p = 0.05. dotted line), throughout the entire range of physiological strain values (0.1 – 0.4) between the perpendicular orientation and the other two orientations, while the difference in shear moduli between the parallel and across orientations did not reach the significance level. The color of the dashed lines represents the two orientations that are being compared from panel A. p-values were calculated at each strain level using a Wald t-statistic.

Shear modulus did not differ significantly between muscle types (Fig. 6). We found no significant effect of including muscle type in our model of shear modulus (F_12,73_ = 1.2, p = 0.33). While EDL and SOL had similar shear moduli throughout the physiological range, the shear modulus of the TA was consistently lower than the other two muscles (Fig. 6A). Though this difference never did reach statistical significance (Fig, 6B), the shear modulus in the TA was as much as 33% and 25% lower than the shear modulus of the EDL and SOL respectively.

**Figure 6.**
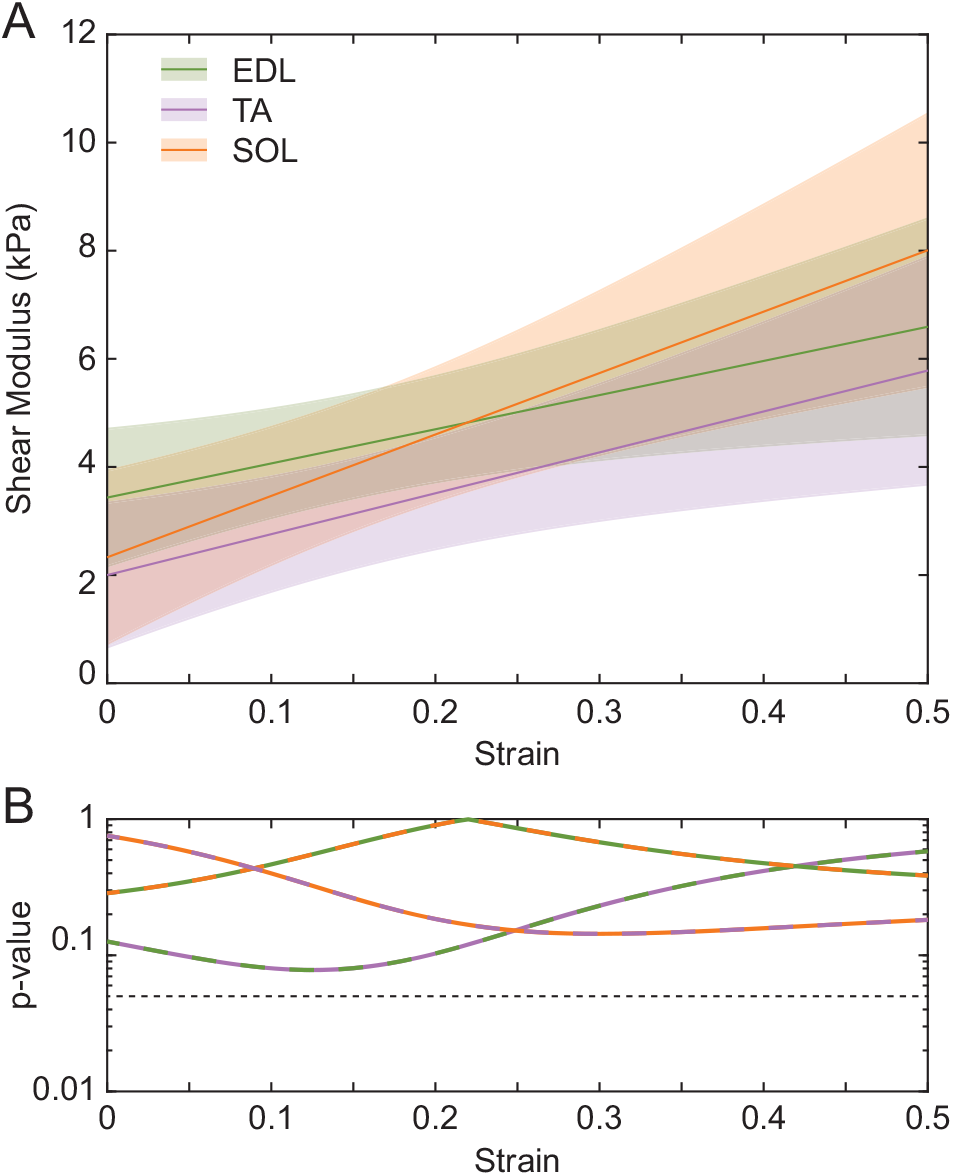
Shear modulus did not differ with muscle architecture. A) Shear modulus was slightly smaller in the TA compared to the EDL and SOL, though any differences were small compared to the uncertainty in the estimates of shear modulus. Shaded area represents 95% confidence interval. B) None of these differences in shear moduli reached the level of significance (p = 0.05. dotted line). The color of the dashed lines represents the two muscles that are being compared from panel A. p-values were calculated at each strain level using a Wald t-statistic.

## Discussion

The objective of this study was to quantify the shear moduli of skeletal muscle in the three orientations relevant to the function of whole muscles. We performed direct mechanical testing on tissue specimens from three muscles of differing architecture in the rat hindlimb. We found that across all muscles and orientations, shear modulus increased linearly with increasing shear strain. We also found that the shear modulus measured perpendicular to the muscle fibers was greater than the shear modulus measured parallel to the fibers or across the fibers. We found no significant difference between the muscles with differing muscle architecture. These results demonstrate the importance in considering the anisotropy of muscle when modeling shear forces in muscles and the role they play in force transmission and muscle injury.

The shear modulus increased linearly with increasing strain. Muscle has been shown to exhibit a nonlinear stress-strain response when subjected to longitudinal extensions (Morrow et al., 2010). We found a similar response to shear deformations (Fig. 3), which resulted in a linear shear modulus-strain relationship (Fig. 4). This relationship is perhaps due to the nonlinear mechanics of the extracellular matrix. In longitudinal extension, muscle fibers and fiber bundles had a linear stress-strain relationship, while the inclusion of extracellular matrix made it nonlinear (Meyer and Lieber, 2011). The extracellular matrix is composed of many wavy collagen fibers (Gillies and Lieber, 2011; Trotter and Purslow, 1992). Therefore, the non-constant shear modulus for muscle, which increases linearly with strain over the full range of physiological values, may be a result of unequal straightening of the extracellular matrix’s collagenous structure (Munster et al., 2013; Sopher et al., 2018). Lengthening of the extracellular matrix due to shearing decreases the number of slack collagen fibers. The addition of more load bearing collagen fibers at greater shear strains may correspond to the increase in shear modulus we observed.

The shear modulus measured perpendicular to fibers was significantly greater than the shear modulus measured parallel to or across fibers. This difference could be due to muscle’s directionally-dependent composition (Gans, 1982; Lieber and Friden, 2000). Muscle is composed of two macroscopic structures, muscle fibers and the extracellular matrix, both of which have unique orientations in muscle (Eng et al., 2008; Purslow and Trotter, 1994; Trotter and Purslow, 1992). It is possible that the muscle fibers have a greater shear modulus than the surrounding extracellular matrix. Therefore, shearing perpendicular to fibers could result in a larger shear modulus because of the resistance caused by the extracellular matrix and muscle fibers, whereas shearing in the parallel and across directions may only have resistance due to the extracellular matrix and transmembrane proteins. This difference could also be in response to how the muscle is physiologically loaded. Under normal physiological conditions, the body does not commonly experience shearing forces across the face of muscle fibers (Blemker et al., 2005; Purslow, 2002). One possible exception to this is blunt trauma where the muscle may be sheared in all three directions (Best, 1997; Loerakker et al., 2013; Woodhouse and McNally, 2011). However, for normal force transmission the structural arrangement of muscle doesn’t provide, or necessitate, a large shear stiffness in the across direction.

Recently, ultrasound elastography has been used to estimate values of shear modulus in muscle. Shear moduli have been shown to relate directly to shear wave speed (Cosgrove et al., 2012; Frulio and Trillaud, 2013). However, due to the heterogeneity in muscle, this relationship may differ from what is expected in other soft tissue (Eby et al., 2013). Using the mechanical estimates of the shear moduli that we made in this study, we can assess whether ultrasound elastography produces feasible estimates of shear moduli. Previous studies in cats (Bernabei et al., 2020), pigs (Eby et al., 2013) and humans (Chernak et al., 2013; Gennisson et al., 2010; Wang et al., 2019) exhibited a shear wave speeds ranging from 2 – 15 m/s. These wave speeds would translate to shear moduli in the range of 4 kPa to 225 kPa. The measures we have made are consistent with the low end of the elastographic measures that were made under passive or unloaded conditions. However, there is quite a large gap between our mechanical measures and the elastographic measures made at high muscle contraction levels. This may be due to shear modulus changing with muscle activation, a phenomenon that cannot be captured in dissected muscle study like ours. Alternatively, it is possible that shear modulus is not the only determinant of shear wave speed (Martin et al., 2018).

### Limitations

The results presented here approximate the material properties of muscle as a uniform material with anisotropic properties. However, muscle is a complex heterogeneous material. The shear moduli we measured are influenced by the constituents of muscle including its extracellular matrix and muscle fibers. Without investigating these materials separately, one cannot decipher their individual contributions. We selected a slow rate of extension to eliminate the viscoelastic effects of muscle from our measures. The rate used is most likely below typical physiological speeds where viscoelastic properties may be important for normal muscle function. Studies at higher rates, capable of measuring the viscoelastic properties, may supplement the measures collected here.

All samples were cut into cubes by hand, which may have introduced slight variations in shape. This was mitigated to the best of our ability by taking dimensional measures before testing and by using a large sample size. Still, this manual approach may have contributed to the large variability in our measures.

## Conclusions

Our results are the first direct measures of three-dimensional shear moduli in skeletal muscle. They show that the shear modulus measured perpendicular to fibers was greater than any other direction.

Additionally, the shear modulus increased linearly with increasing strain, indicative of a nonlinear elastic material. Despite architectural differences between muscles, we did not find the shear modulus of the rat muscles we tested to be different. The quantitative measures reported here can be used to describe the mechanical properties of muscle in shear, which adds to the growing knowledge of muscle material properties as are needed to develop computational models of how muscle responds to the complex stresses and strains essential to its use in the control of movement and posture.

## Conflict of Interest Statement

The authors have no financial or personal conflicts of interest to disclose.

## Acknowledgments

This work was supported by National Institute of Health grants R01AR071162 and T32HD07418.

## Notes

### Competing Interest Statement

The authors have declared no competing interest.

## References

Bernabei, M., Lee, S.S.M., Perreault, E.J., Sandercock, T.G., 2020. Shear wave velocity is sensitive to changes in muscle stiffness that occur independently from changes in force. J Appl Physiol (1985) 128, 8–16.

Best, T.M., 1997. Soft-Tissue Injuries and Muscle Tears. Clinics in Sports Medicine 16, 419–434.

Blemker, S.S., Delp, S.L., 2005. Three-dimensional representation of complex muscle architectures and geometries. Ann Biomed Eng 33, 661–673.

Blemker, S.S., Pinsky, P.M., Delp, S.L., 2005. A 3D model of muscle reveals the causes of nonuniform strains in the biceps brachii. Journal of Biomechanics 38, 657–665.

Bosboom, E.M.H., Hesselink, M.K.C., Oomens, C.W.J., Bouten, C.V.C., Drost, M.R., Baaijens, F.P.T., 2001. Passive transverse mechanical properties of skeletal muscle under in vivo compression. Journal of Biomechanics 34, 1365–1368.

Burkholder, T.J., Fingado, B., Baron, S., Lieber, R.L., 1994. Relationship between Muscle-Fiber Types and Sizes and Muscle Architectural Properties in the Mouse Hindlimb. Journal of Morphology 221, 177–190.

Chernak, L.A., DeWall, R.J., Lee, K.S., Thelen, D.G., 2013. Length and activation dependent variations in muscle shear wave speed. Physiol. Meas. 34, 713–721.

Cosgrove, D.O., Berg, W.A., Dore, C.J., Skyba, D.M., Henry, J.P., Gay, J., Cohen-Bacrie, C., Group, B.E.S., 2012. Shear wave elastography for breast masses is highly reproducible. Eur Radiol 22, 1023–1032.

Eby, S.F., Song, P., Chen, S., Chen, Q., Greenleaf, J.F., An, K.N., 2013. Validation of shear wave elastography in skeletal muscle. J. Biomech. 46, 2381–2387.

Eng, C.M., Smallwood, L.H., Rainiero, M.P., Lahey, M., Ward, S.R., Lieber, R.L., 2008. Scaling of muscle architecture and fiber types in the rat hindlimb. J Exp Biol 211, 2336–2345.

Frulio, N., Trillaud, H., 2013. Ultrasound elastography in liver. Diagn Interv Imaging 94, 515–534.

Gans, C., 1982. Fiber architecture and muscle function. Exerc Sport Sci Rev 10, 160–207.

Gennisson, J.L., Deffieux, T., Mace, E., Montaldo, G., Fink, M., Tanter, M., 2010. Viscoelastic and anisotropic mechanical properties of in vivo muscle tissue assessed by supersonic shear imaging. Ultrasound Med. Biol. 36, 789–801.

Gillies, A.R., Lieber, R.L., 2011. Structure and function of the skeletal muscle extracellular matrix. Muscle Nerve 44, 318–331.

Herbert, R.D., Gandevia, S.C., 2019. The passive mechanical properties of muscle. J Appl Physiol (1985) 126, 1442–1444.

Huijing, P.A., 1999. Muscle as a collagen fiber reinforced composite: a review of force transmission in muscle and whole limb. Journal of Biomechanics 32, 329–345.

Jarvinen, T.A., Jarvinen, M., Kalimo, H., 2013. Regeneration of injured skeletal muscle after the injury. Muscles Ligaments Tendons J 3, 337–345.

Lieber, R.L., Friden, J., 2000. Functional and clinical significance of skeletal muscle architecture. Muscle Nerve 23, 1647–1666.

Lieber, R.L., Friden, J., 2019. Muscle contracture and passive mechanics in cerebral palsy. J Appl Physiol (1985) 126, 1492–1501.

Loerakker, S., Solis, L.R., Bader, D.L., Baaijens, F.P., Mushahwar, V.K., Oomens, C.W., 2013. How does muscle stiffness affect the internal deformations within the soft tissue layers of the buttocks under constant loading? Comput Methods Biomech Biomed Engin 16, 520–529.

Luke, S.G., 2017. Evaluating significance in linear mixed-effects models in R. Behav Res Methods 49, 1494–1502.

Martin, J.A., Brandon, S.C.E., Keuler, E.M., Hermus, J.R., Ehlers, A.C., Segalman, D.J., Allen, M.S., Thelen, D.G., 2018. Gauging force by tapping tendons. Nat Commun 9, 1592.

Meyer, G.A., Lieber, R.L., 2011. Elucidation of extracellular matrix mechanics from muscle fibers and fiber bundles. Journal of Biomechanics 44, 771–773.

Morrow, D.A., Haut Donahue, T.L., Odegard, G.M., Kaufman, K.R., 2010. Transversely isotropic tensile material properties of skeletal muscle tissue. Journal of the Mechanical Behavior of Biomedical Materials 3, 124–129.

Munster, S., Jawerth, L.M., Leslie, B.A., Weitz, J.I., Fabry, B., Weitz, D.A., 2013. Strain history dependence of the nonlinear stress response of fibrin and collagen networks. Proc Natl Acad Sci U S A 110, 12197–12202.

Purslow, P.P., 2002. The structure and functional significance of variations in the connective tissue within muscle. Comparative Biochemistry and Physiology Part A: Molecular & Integrative Physiology 133, 947–966.

Purslow, P.P., Trotter, J.A., 1994. The morphology and mechanical properties of endomysium in series-fibred muscles: variations with muscle length. Journal of Muscle Research and Cell Motility 15, 299–308.

Sopher, R.S., Tokash, H., Natan, S., Sharabi, M., Shelah, O., Tchaicheeyan, O., Lesman, A., 2018. Nonlinear Elasticity of the ECM Fibers Facilitates Efficient Intercellular Communication. Biophysical journal 115, 1357–1370.

Trotter, J.A., Purslow, P.P., 1992. Functional morphology of the endomysium in series fibered muscles. Journal of Morphology 212, 109–122.

Tuttle, L.J., Alperin, M., Lieber, R.L., 2014. Post-mortem timing of skeletal muscle biochemical and mechanical degradation. Journal of Biomechanics 47, 1506–1509.

Wang, A.B., Perreault, E.J., Royston, T.J., Lee, S.S.M., 2019. Changes in shear wave propagation within skeletal muscle during active and passive force generation. J Biomech 94, 115–122.

Woodhouse, J.B., McNally, E.G., 2011. Ultrasound of skeletal muscle injury: an update. Semin Ultrasound CT MR 32, 91–100.

